# Task-Modulated Neural Responses in Scene-Selective Regions of the Human Brain

**DOI:** 10.1101/2023.12.13.571229

**Authors:** Aysu Nur Koc, Burcu A. Urgen, Yasemin Afacan

## Abstract

The study of scene perception is crucial to the understanding of how one interprets and interacts with their environment, and how the environment impacts various cognitive functions. The literature so far has mainly focused on the impact of low-level and categorical properties of scenes and how they are represented in the scene-selective regions in the brain, PPA, RSC, and OPA. However, higher-level scene perception and the impact of behavioral goals has not been explored in depth. Moreover, the selection of the stimuli has not been systematic and mainly focused on outdoor environments. In this fMRI experiment, we adopted multiple behavioral tasks, selected real-life indoor stimuli with a systematic categorization approach, and used various multivariate analysis techniques to explain the neural modulation of scene perception in the scene-selective regions of the human brain. Participants (N=21) performed categorization and approach-avoidance tasks during fMRI scans while they were viewing scenes from built environment categories based on different affordances ((i)access and (ii)circulation elements, (iii)restrooms and (iv)eating/seating areas). Searchlight-based classification analysis revealed that the OPA was significantly successful in decoding scene category regardless of the task, and that the task condition affected category decoding performances of all the scene-selective regions. Model-based representational similarity analysis (RSA) revealed that the activity patterns in scene-selective regions are best explained by task. These results contribute to the literature by extending the task and stimulus content of scene perception research, and uncovering the impact of behavioral goals on the scene-selective regions of the brain.

## 1. Introduction

A scene is often defined as a view of an environment with a spatial structure one can act within (R. Epstein et al., 1999; R. Epstein, 2005; Henderson & Hollingworth, 1999). The human brain can process incredibly large amounts of data regarding the environment one is situated within in a matter of milliseconds, which is often attributed to various evolutionary mechanisms, such as identifying one’s surroundings, finding shelter, food, and protection from threats (Kaplan, 1992). Providing the background and the context for virtually any cognitive process, the study of scene perception can be crucial for fully understanding the human brain. It can contribute to the study of many topics, such as attention, learning, memory, and social behavior (R. A. Epstein & Baker, 2019). Accordingly, this significance of scenes has inspired a large body of research investigating the behavioral and neural bases of scene perception and their implications.

Three main regions were discovered to be selective to scenes in the human brain compared to other categorical stimuli. The parahippocampal place area (PPA) is located in the posterior parahippocampal gyrus and is highly sensitive to scenes; it shows increased activation to stimuli with spatial layout, including photos, drawings, and even spaces built with Lego blocks (R. Epstein et al., 1999; R. Epstein & Kanwisher, 1998; Walther et al., 2011). PPA was demonstrated to be sensitive to spatial, categorical, and semantic properties of scenes that help us recognize scenes and distinguish between them (Persichetti & Dilks, 2018, 2019). In contrast, it was not emphasized in processes related to navigation, planning, or memory regarding scenes.

The retrosplenial cortex (RSC) is located in the posterior cingulate region (Brodmann’s 29 and 30). In addition to its scene-selectivity, it is emphasized in processes such as learning, navigation, and episodic memory regarding scenes (Vann et al., 2009). In contrast, it may not be as successful as the other regions regarding scene category sensitivity (Persichetti & Dilks, 2019). Instead, it is viewed as an association hub integrating a wide range of information regarding one’s environment (Alexander et al., 2022), necessary for higher-level complex processes such as map-based navigation and route-planning (Dilks et al., 2022), supported by its sensitivity to heading direction, reference frames, boundaries, landmarks (Alexander et al., 2022; Auger et al., 2012; Dilks et al., 2011; Stacho & Manahan-Vaughan, 2022; Troiani et al., 2014), and role in reconstructing places by prediction or by retrieval from memory (R. A. Epstein, Higgins, et al., 2007; R. A. Epstein, Parker, et al., 2007).

Lastly, the occipital place area (OPA) is located near the transverse occipital sulcus, and its activation is modulated by egocentric distance, first-person perspective motion, direction, obstacles, and borders in a scene (Dilks et al., 2022; Henriksson et al., 2019; Kamps et al., 2016). Considering these findings, the OPA is thought to have a major role in perceiving and navigating the immediate scene one is situated within (called *visually guided navigation*) rather than complex navigation referring to memory or experience (Dilks et al., 2013; Silson et al., 2015). Its disruption can cause problems in scene categorization (Ganaden et al., 2013); however, this is thought to be related to the OPA’s sensitivity to border and layout that could differentiate scenes and not to semantic processing like the PPA. In addition to its role in visually-guided navigation, the OPA is suggested to be an information source for higher processing of scenes in the PPA and RSC (Dilks et al., 2013).

These scene-selective regions and the underlying mechanisms of scene perception have been studied using various stimuli and tasks, contributing to our current knowledge. Scene perception research often employs categorical stimuli. Common categories of scene stimuli include various outdoor categories such as mountains, fields, and beaches (M. Greene & Oliva, 2009; Johnson & Johnson, 2014), indoors and outdoors (M. R. Greene & Hansen, 2020; Henderson et al., 2007; Rousselet et al., 2005), natural and manmade environments (Gopnarayan et al., 2022; Groen et al., 2013), environments that vary by their openness and closedness (Guo et al., 2012; Kravitz et al., 2011), or by other distinctions including properties such as scale, spatial frequency, and color (Kaping et al., 2007; Oliva & Schyns, 2000; Oliva & Torralba, 2001). These categorical stimuli are often presented to the participants with an accompanying task, such as categorization to measure their understanding of the stimuli, or tasks that aim to keep the participants focused on the screen, such as n-back paradigms.

Although past research uncovered so much about scene processing dynamics in the human brain, there are several gaps in knowledge, especially regarding higher-level scene perception. First, the field lacks the use of valid behavioral tasks that are related to goals one can have regarding these scenes and were limited to tasks that purely aim to keep the participant focused (excluding research specifically focusing on navigation and memory, which is beyond the scope of this paper). The effect of task on brain activity patterns while participants look at the same stimuli has been shown for other category-sensitive brain regions (Harel et al., 2014); however, such comparisons are limited for scene-selective regions. Given the growing interest in the role of tasks in visual neuroscience (Kay et al., 2023), it has become important to study how the neural representations of scenes change under different task demands. Second, the approach to choosing stimulus categories has not been systematic, and the stimulus content has been limited. Categories of stimuli used in past research are primarily outdoor scenes, and categorical distinctions are decided on based seemingly on common sense.

Further, built indoor spaces in which we spend 80 to 90% of our daily lives (Cholowsky et al., 2023; Klepeis et al., 2001) and made up of highly functional parts that are suitable for studying various behavioral goals are largely overlooked. Moreover, the stimuli representing these categories are often professional high-quality images, previously referred to by Epstein et al. as “holiday snap-shot perception” (R. A. Epstein & Baker, 2019) or the complete opposite: images stripped off of their visual properties, reducing them to a singular aspect a particular study focuses on, neither of which reflecting real-life scenes we interact with (Groen et al., 2017). This need for ecologically valid behavioral tasks and stimuli in the scene literature for a more comprehensive and high-level study of scenes both behaviorally and neurally has been pointed out multiple times in the past (R. A. Epstein & Baker, 2019; Groen et al., 2017; Malcolm et al., 2016).

In this study, we aim to address these gaps by examining how different behavioral goals modulate the neural processing of built environment categories in the scene-selective regions of the human brain. To this end, we employed two behavioral tasks in an fMRI experiment while participants viewed indoor scenes: one is a categorization task to measure the semantic processing of categories and monitor the participants’ understanding of categories, and the other is an approach-avoidance task measuring initial actions taken by participants regarding a scene. We studied these tasks in less studied indoor environments, comprising separate functional units suitable for studying behavioral goals. To systematically decide on indoor categories, we adapted a categorization method from the architecture literature, defining built-environment categories based on their elements that afford distinct actions and serve distinct functions. Finally, we selected real-life scene stimuli from a database to keep the stimuli natural and ordinary, reflecting real-life conditions. We then used multivariate pattern analysis (MVPA) and representational similarity analysis (RSA) to examine how tasks modulate the neural representation of scenes in the scene-selective regions of the human brain.

## 2. Methods

### 2.1 Participants

28 participants volunteered for this fMRI experiment. After data quality checks explained in section 2.6, the final analyses included 21 participants (11 males, ages 18-31, M=23.7, SD=3.25). All participants had normal or corrected-to-normal visual acuity, reported no neurological disorders, and did not use any related medication. The study was approved by the Human Research Ethics Committee of Bilkent University in line with the Declaration of Helsinki. Each participant filled out a prescreening form for MR safety and gave written consent before the experiment. After the experiment, they were briefed upon request and compensated for their time, with course credit when applicable and 100₺.

### 2.2 Stimuli and Apparatus

Stimulus categories were chosen based on the Universal Design: Users-Built Environments Model by Froyen (2012), which evaluates built environments from various aspects and provides an in-depth classification. The model dissects environments based on the impairments and activities of users, the physical aspects such as lighting and ergonomics, and the physical concrete elements. The *elements* section of the model separates environments into physical parts: approach (the surrounding), access (entrances), horizontal circulation (e.g., corridors), vertical circulation (e.g., stairs), social interaction (e.g., canteens), rest (waiting and seating areas), food and drinks (food courts), and sanitary facilities (restrooms). Considering the objectives of this study, we selected only the elements section of this categorization, as these were less subjective, and they categorized environments based on more concrete and visually distinct elements. However, since some of these categories had shared features, we grouped them and created a simplified, easier-to-understand categorization approach. The final set of experimental stimuli consisted of 32 different images chosen from the Scene Understanding (SUN) database (Xiao et al., 2010, 2016). There were 8 images of access elements to buildings (4 entrances, 4 exit points), 8 images of circulation elements (2 stairs, 2 escalators, 2 corridors, and 2 elevators), 8 images of sanitary facilities (4 bathroom stalls, 4 sink areas), and 8 images of seating areas (4 eating areas, 4 seating areas). Simplified categories and the corresponding stimuli used in the experiment can be seen in Fig. 1.

**Fig. 1.**
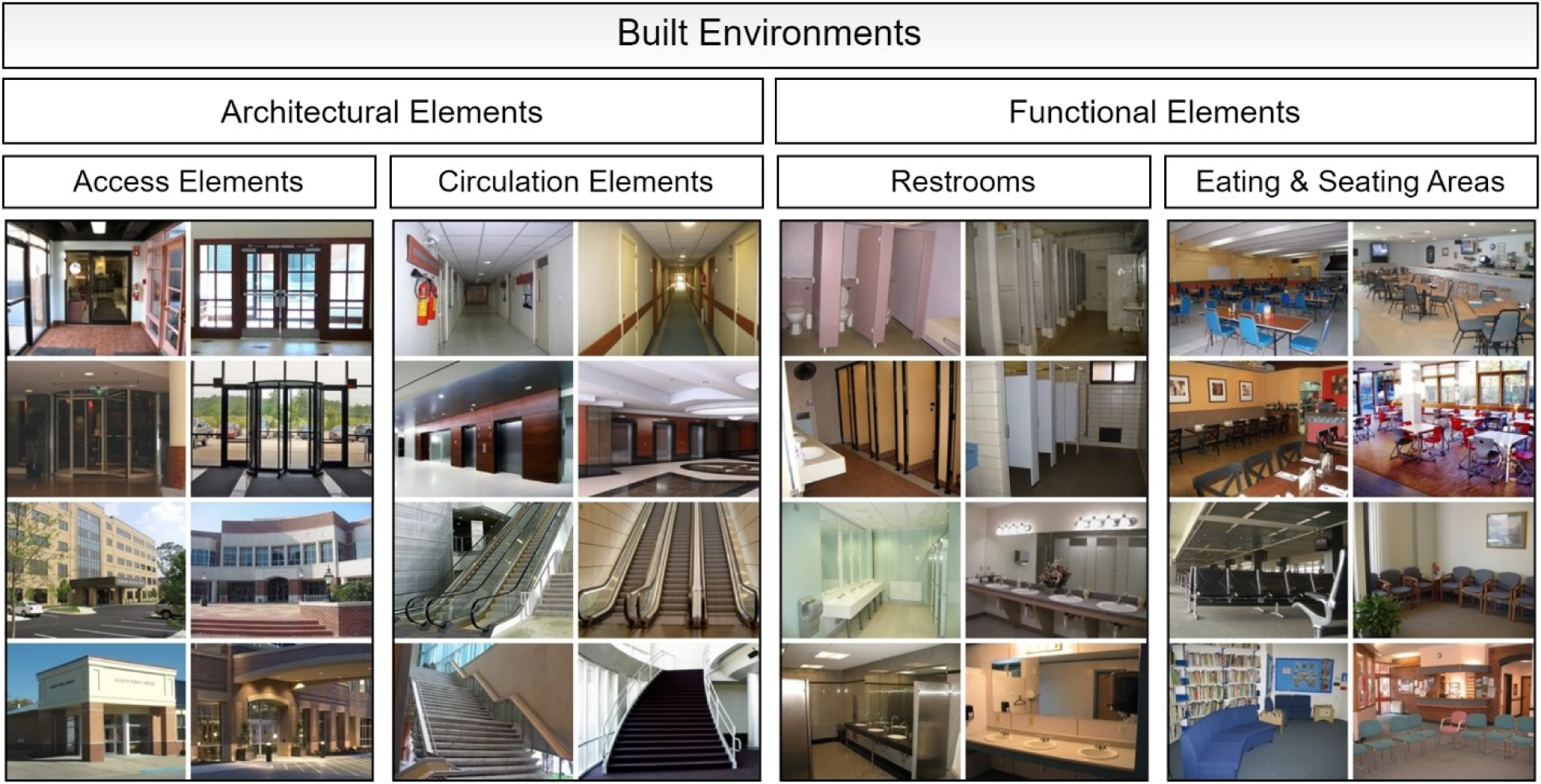
Simplified stimulus categories and the corresponding stimuli.

Stimuli were presented on the screen with a 600×450 resolution. The visual properties of images were not manipulated to keep the stimuli ecologically valid, but these properties were addressed in the analyses.

### 2.3 Behavioral Tasks

Participants performed 2 different tasks during the experiment. One of these was a categorization task, where participants responded by indicating the main category of the presented stimulus (architectural or functional). The categories required considering different actions a scene affords, along with understanding its semantic associations, so this task aimed to engage participants with various high-level processes needed to comprehend a scene wholly.

The other was an approach-avoidance task, where they simply indicated whether they would like to enter the presented environment. We address this task as an “approach-avoidance task”; however, it must be emphasized that it does not precisely represent the nature of such tasks found in the literature. Approach-avoidance tasks often involve distinctly separated stimuli based on a positive-negative dynamic related to valence (Korn & Elliot, 2015). Therefore, it is more sensible to consider this task a subjective “enter-or-not” decision since the stimuli were not chosen based on valence. Instead, we aim to observe how an immediate decision about acting in an environment is made, that is, to enter or not, based on initial perception. Previous research in neuroarchitecture literature shows that approach-avoidance decisions are not entirely explained by beauty or otherwise valence-based responses and that these tasks can measure initial sensorimotor processing regarding an environment (Coburn et al., 2020). Participants were also instructed with this in mind: they were asked to respond quickly based on their initial opinions. They were not expected to be consistent across runs for the same stimuli. Since there were no correct answers, this task was not analyzed or interpreted like a traditional approach-avoidance task and is better considered unique to this study.

### 2.4 Experimental Design

The experiment consisted of 8 runs (∼9 minutes each), containing 2 behavioral task blocks (categorization and approach-avoidance) with their order counterbalanced across runs (ABBABAAB). With one trial per stimulus, each task consisted of 32 trials. Trials began with a 2000ms presentation of the stimulus, followed by a 2000ms response period, and ended with a randomized variable inter-stimulus interval (ISI) that ranged between 3000-4000ms, averaging at 3500ms. Each task block started with a 10-second instruction screen indicating the task and the corresponding buttons and ended with a 10-second rest. A black fixation cross was presented at the center of the screen throughout the experiment, which only turned white during the response period. The procedure of a single run is shown in Fig. 2.

**Fig. 2.**
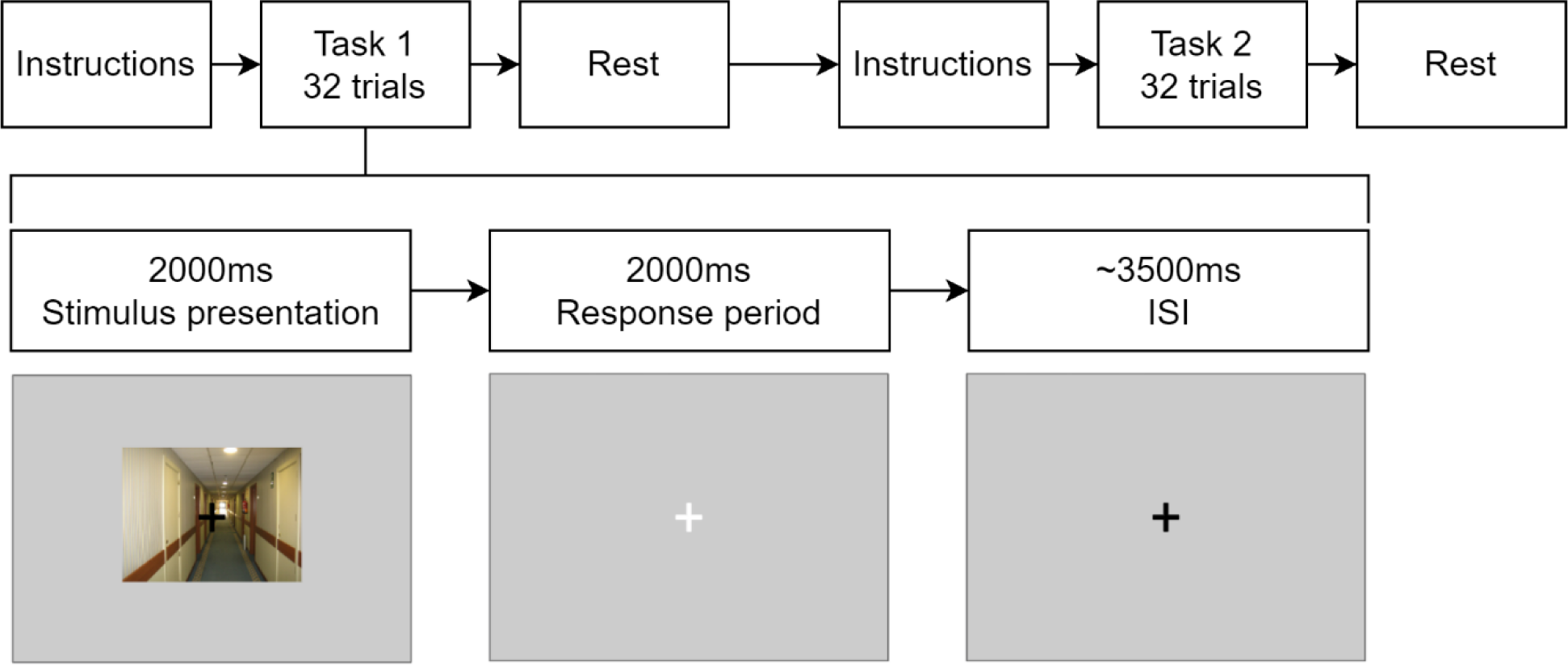
Experimental procedure of a run. Each run consists of two task blocks, each including 32 trials with 2000ms stimulus presentation, 2000ms response period, and ∼3500ms ISI.

The categories were explained to the participants in detail, and they also performed a shorter version of these tasks on a laptop as a practice task before going into the MRI scanner. Architectural elements comprising access points and circulation elements were explained to the participants as parts of buildings that allow us to access and circulate within an environment, and functional elements comprising restrooms and eating/seating areas were explained as those that serve basic human needs and comprise relatively stationary activities. Participants were instructed to focus on the fixation throughout the experiment. The responses were collected using a fiber optic response box.

### 2.5 Functional Localizer

In addition to the experimental session, a separate functional localizer session was conducted to delineate scene-selective regions of interest (ROIs) across our participants. This localizer was adapted from the code made available by Meissner et al. (Meissner et al., 2019) and was shown to activate scene-selective regions in multiple age groups successfully. This session consisted of 4 short runs (∼3.5 minutes) of scene, object, and fixation blocks presented rapidly while participants performed a one-back repetition task.

### 2.6 FMRI Data Acquisition and Preprocessing

MRI data was collected using a 3T Siemens Trio scanner (Magnetom Trio, Siemens AG, Erlangen, Germany) with a 32-channel head coil, at the National Magnetic Resonance Research Center (UMRAM), İhsan Doğramacı Bilkent University. Participants viewed visual stimuli presented on an MRI-safe LCD screen (1920×1080px, 125×70cm, vertical refresh rate=60Hz, TROYKA MED, İstanbul) through a mirror mounted on the head coil with a total view distance of ∼168cm. The sessions started with a short localizer scan to check if the head was positioned correctly, followed by a T1-weighted structural scan. The main experimental session included 8 functional runs, each lasting about 9 minutes. The high resolution T1-weighted structural images were obtained with a standard protocol (TR=2600ms, TE=2.92ms, flip angle=12°, FoV read=256mm, FoV phase=87.5%, 176 slices, voxel size=1×1×1mm3). During the functional runs, 263 functional volumes were obtained using gradient-echo planar imaging (TR=2000ms, TE=22ms, flip angle=90°, 64×64 matrix, FoV read=192mm, 43 slices with a thickness of 2.5mm, voxel size=3×3×2.5mm3). For the localizer session for ROI definition, the fMRI data acquisition parameters were the same as the main experiment, only this time, we obtained 4 functional runs (∼3.5 minutes/104 volumes)

Anatomical and functional fMRI data was converted to BIDS format (Gorgolewski et al., 2016) and later preprocessed using fMRIPrep 21.0.1 (Esteban, Blair, et al., 2018; Esteban, Markiewicz, et al., 2018) using the default steps. Structural (T1w) images were corrected for intensity non-uniformity and skull stripped. Brain tissue segmentation was performed, and brain surfaces were reconstructed. Volume-based spatial normalization to the ICBM 152 Nonlinear Asymmetrical template (2009c) was performed. For the functional images (both the main experiment and the functional localizer), a reference volume and its skull-stripped version were created, and head-motion parameters were calculated based on the reference. Slice-time correction was applied to BOLD runs and then resampled onto the native space after head-motion estimation transforms were applied. The BOLD reference was co-registered to the T1w reference with six degrees of freedom. Confounding time-series were calculated for framewise displacement, demeaned variance (DVARS), global signals, and physiological regressors to be used for component-based noise correction. High-pass filtering was applied to the preprocessed BOLD time-series with a 128s cut-off. BOLD time-series were also resampled into the same standard space as the structural scans. Detailed preprocessing documentation outputted by the software itself can be found in the supplementary material.

Based on the head-motion output of this procedure, runs with spikes of motion of more than 1.5mm and total relative motion of more than 3mm were defined and removed from the analysis. Participants with less than 5 suitable runs remaining and those who did not attend the localizer session were completely omitted from the analyses. Therefore, the final analyses were performed on 21 participants in total, with complete data (8 runs) from 17 participants, 7 runs from 1, 6 runs from 2, and 5 runs from 1 participant.

### 2.7 Data Analyses

#### 2.7.1 Behavioral Analyses

The accuracy of responses was calculated for the categorization task. Since there were no correct responses for the approach-avoidance task, only the distribution of responses was examined. Reaction times were compared across tasks using a paired t-test.

#### 2.7.2 Functional Localizer and Definition of Regions of Interest (ROIs)

Functional localizer scans (and also the GLM analyses in the following sections) were analyzed using the Statistical Parametric Mapping software package (SPM12, Wellcome Trust Centre for Neuroimaging, University College London, UK) implemented under MATLAB (The Mathworks Inc., Natick, MA). Preprocessed functional scans were smoothed with a 6mm full width at half maximum (FWHM) kernel. The general linear model (GLM) was calculated using a design matrix consisting of 11 regressors, including scene, object, and fixation blocks, as well as the rest of the intervals in the experimental procedure and 6 motion regressors. Scenes>objects contrast images were created for each participant. One sample t-test was performed at the group level using these contrast images with default SPM parameters.

#### 2.7.3 Searchlight-based Multivariate Pattern Analysis (MVPA)

We performed a searchlight-based classification analysis to observe classification performance for tasks and categories in these ROIs. First, we performed GLM analyses on the unsmoothed experimental functional scans using SPM12. Contrast images were created by grouping trials for each category under each task (4 categories during the categorization task and 4 during the approach-avoidance task).

For the classification analysis, we used The Decoding Toolbox (TDT - version 3.999F) (Hebart et al., 2015), implementing a linear support vector machine (LibSVM) (Chang & Lin, 2011) with a fixed cost parameter c=1. Classifiers were trained with a leave-one-run-out procedure using runwise beta images of participants and repeated multiple times to test for each run while training with the rest (k=8 for 17, k=7 for 1, k=6 for 2, and k=5 for 1 participant). We moved a 4-voxel searchlight within the bilateral ROI masks for each participant. This procedure was performed 4 times in total, one decoding for categories (labels: 1,2,3,4,1,2,3,4), decoding for the task (1,1,1,1,2,2,2,2), for categories during the categorization task (1,2,3,4,0,0,0,0), and lastly for categories during the approach-avoidance task (0,0,0,0,1,2,3,4). For each classification, accuracy-minus-chance images and average accuracy values were computed for participants individually. One-way Analysis of Variance (ANOVA) was performed separately to analyze differences in category and task decoding performances across ROIs, and a two-way ANOVA was performed for the categorization performance of ROIs across tasks (2 tasks x 3 ROIs). Additionally, one-sample t-tests were applied for each decoding condition to each ROI to determine if the decoding performances were significantly higher than the chance level.

#### 2.7.4 Representational Similarity Analysis (RSA)

To examine the changes in activation patterns for each condition across ROIs compared to tasks, categories, and visual image properties, we performed a model-based representational similarity analysis (RSA) using the RSA Toolbox (Nili et al., 2014). RSA allows us to explore how information is represented in the brain by comparing patterns of neural activity to different measurements or conditions, such as stimuli, tasks, and hypothetical models. By calculating representational dissimilarity matrices (RDMs) of neural activity in different brain regions and comparing them to matrices of other measurements or models, it helps discern the underlying structure of neural activities (Kriegeskorte, 2008).

First, we created 4 different 64×64 candidate models that could potentially explain the changes in activation patterns in the form of RDMs (Fig. 3). The first was a task model, indicating that the dissimilarity values would be low for all conditions during the same task and high during the other. The second was a category model with low dissimilarity values between items of the same category and higher dissimilarity between items of other categories independent of the task. The third was a visual model created using Gabor filters on the stimuli to account for the visual differences among the stimuli, as scene processing is affected and sometimes explained by low-level properties (Groen et al., 2017; Kauffmann et al., 2014). A Gabor bank was defined with 3 wavelengths (4,8,16) and 4 orientations (0,45,90,135). Gabor filters of each combination were applied to the stimuli, and the resulting magnitude information was converted into arrays and combined for each of the 32 stimuli. Later, the resulting 32 arrays were compared to one another (corr2 in MATLAB). The obtained similarity values were subtracted from 1 to calculate dissimilarity values and placed in a 32×32 matrix. This matrix was repeated 2×2 to create a 64×64 model. The fourth and final model was a random one for control.

**Fig. 3.**
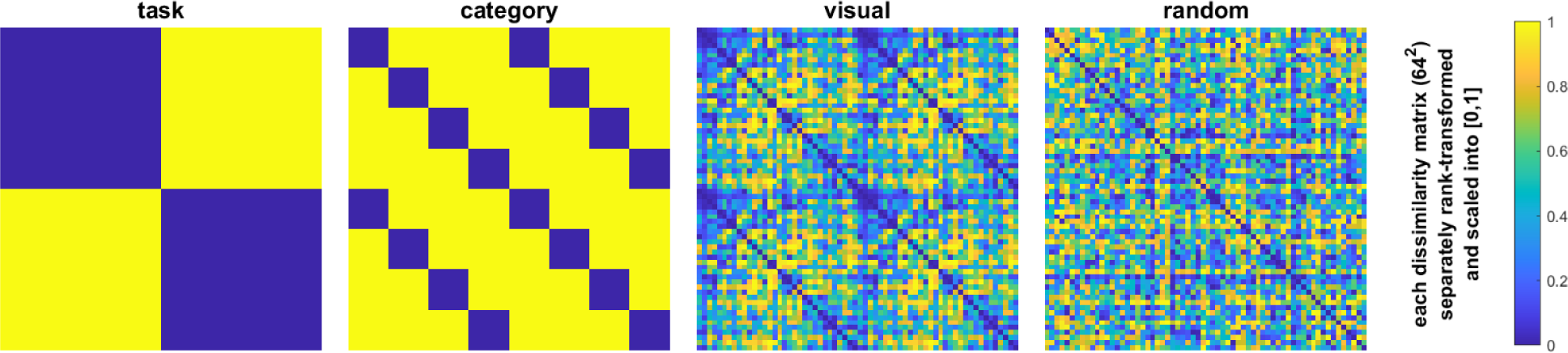
Candidate model RDMs created for the RSA.

Later, we performed another GLM analysis on the unsmoothed data, with the only difference from the previous sections being that we defined each stimulus under each task as a separate condition this time, resulting in 64 conditions. The obtained thresholded images (spmT) were used as input for the RSA Toolbox. Averaged 64×64 RDMs were computed across 21 subjects. Then, via Kendall’s τ, a one-sided Wilcoxon signed-rank random-effects test was performed between the neural RDMs and the candidate models explained above. By estimating a hypothetical model with maximum average correlation to the reference RDM via Kendall’s τ, the upper and lower bounds for the noise ceiling were calculated, indicating the range a model must reach to be considered a *true model* fully explaining the variance in the data. Lastly, by calculating Spearman’s ρ, pairwise correlations of all the neural and model RDMs were compared. Both for ROI to candidate model comparisons and overall RDM comparisons, multidimensional scaling (MDS) plots were also generated. Matrix values of both model and neural RDMs were rank-transformed into [0,1] and color-coded accordingly.

## 3. Results

### 3.1 Behavioral Results

Participants responded accurately 97.014% of the time during the categorization task, indicating the categorization method was well understood. The reaction times were significantly different between the tasks (p<0.05), with a moderate effect size (Cohen’s d=-0.603).

The distribution of answers for the approach-avoidance task by stimulus and participant can be seen in supplementary figures (Sup. Fig. 1 & 2).

### 3.2 Functional Localizer Results and ROIs

To decide on the maximum extent of our ROIs, we used the scene parcels from a localizer study by Julian et al. with a large participant group (Julian et al., 2012). Group-level activation maps were overlaid by these parcels, and by adjusting the threshold, the strongest responding 50-voxel clusters were selected in each hemisphere at the sites indicated for PPA, RSC, and OPA. Group-level ROIs were defined using the MarsBar toolbox (Brett et al., 2010) by combining these bilateral clusters, resulting in 3 100-voxel ROI masks (Fig. 4). Statistical information regarding these ROIs is listed in Table 1. Peak locations were comparable to the past literature (e.g. Marchette et al., 2015; Ramanoël et al., 2019).

**Fig. 4.**
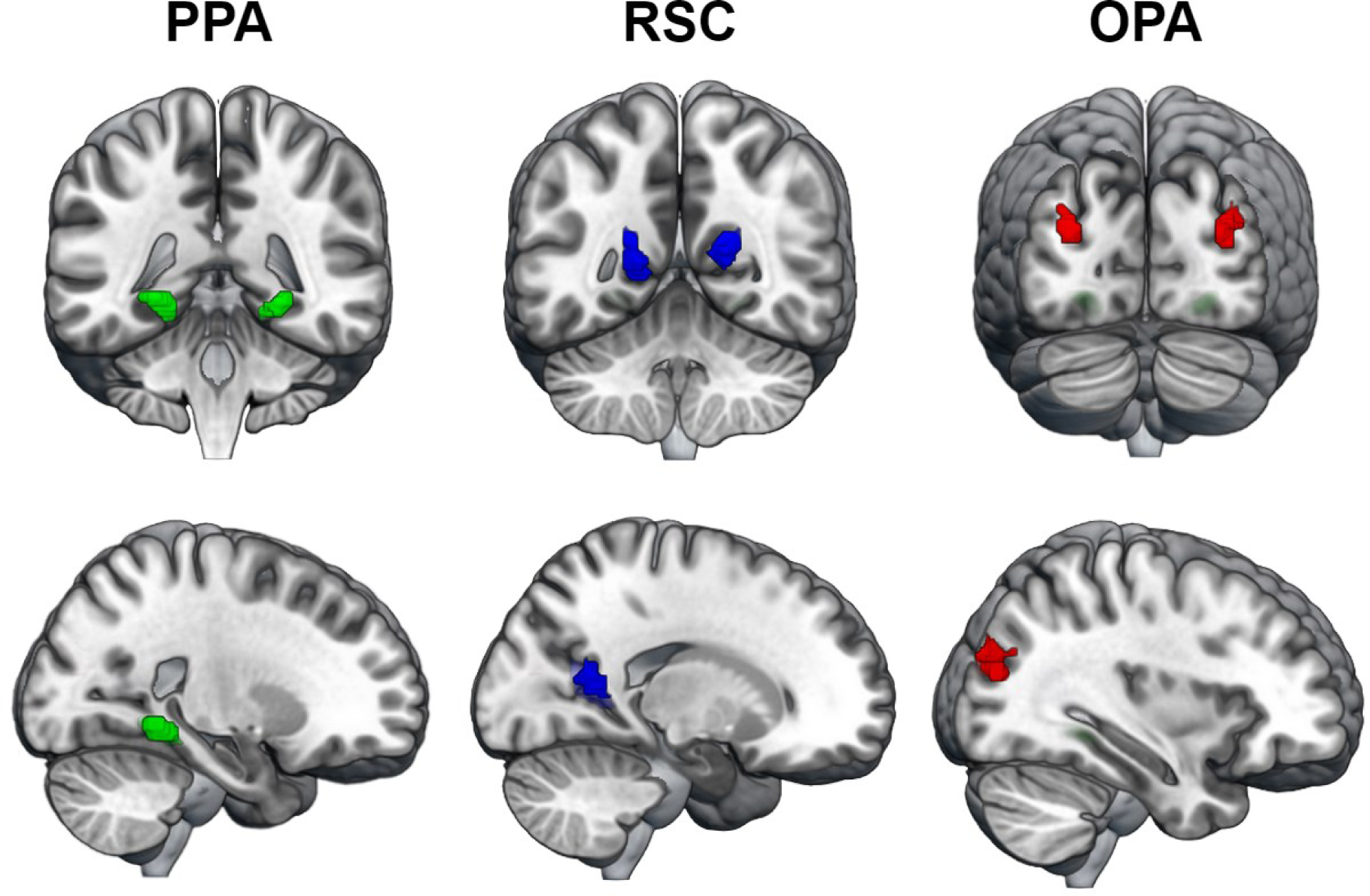
100-voxel bilateral ROI masks, obtained by the group-level results of the localizer session.

**Table 1.** Coordinates and peaks of the ROI clusters.

| ROI | Left Hemisphere |  |  | Right Hemisphere |  |  |
| --- | --- | --- | --- | --- | --- | --- |
|  | Cluster Threshold | Peak Threshold | Peak Coordinates | Cluster Threshold | Peak Threshold | Peak Coordinates |
| PPA | T=9 | T=15.42<br>T=9.41 | -24 -48 -6<br>-22 -36 -14 | T=9.2 | T=19.71 | 26 -46 -6 |
| RSC | T=6.5 | T=11.27<br>T=8.43<br>T=5.43 | -18 -58 9<br>-16 -48 4<br>-6 -42 2 | T=8.5 | T=14.11 | 20 -54 14 |
| OPA | T=3.95 | T=6.57<br>T=5.05 | -30 -84 22<br>-34 -82 32 | T=6.06 | T=8.25<br>T=8.07<br>T=6.72 | 32 -78 19<br>32 -84 26<br>38 -78 26 |

### 3.3 Searchlight-based MVPA Results

The findings related to the decoding performance differences of ROIs by task, category, and category decoding tasks can be seen in Fig. 5. One-way ANOVA analyzing the task decoding performances of ROIs revealed no significant effects (F(2,60)=0.52, p=0.598). Another one-way ANOVA was performed to investigate the category decoding performances of ROIs, and it revealed statistically significant effects (F(2,60)=4.69, p=0.013). Post-hoc tests with Bonferroni correction revealed a significant difference between the RSC and the OPA, at p=0.014. Finally, a two-way ANOVA was performed to analyze the effect of tasks on category decoding performance across ROIs. This analysis did not reveal any effect of ROI (p=0.845) or task (p=0.976) on the category decoding performance, nor any significant interaction between ROI and task (F(2,120)=0.47, p=0.627).

**Fig. 5.**
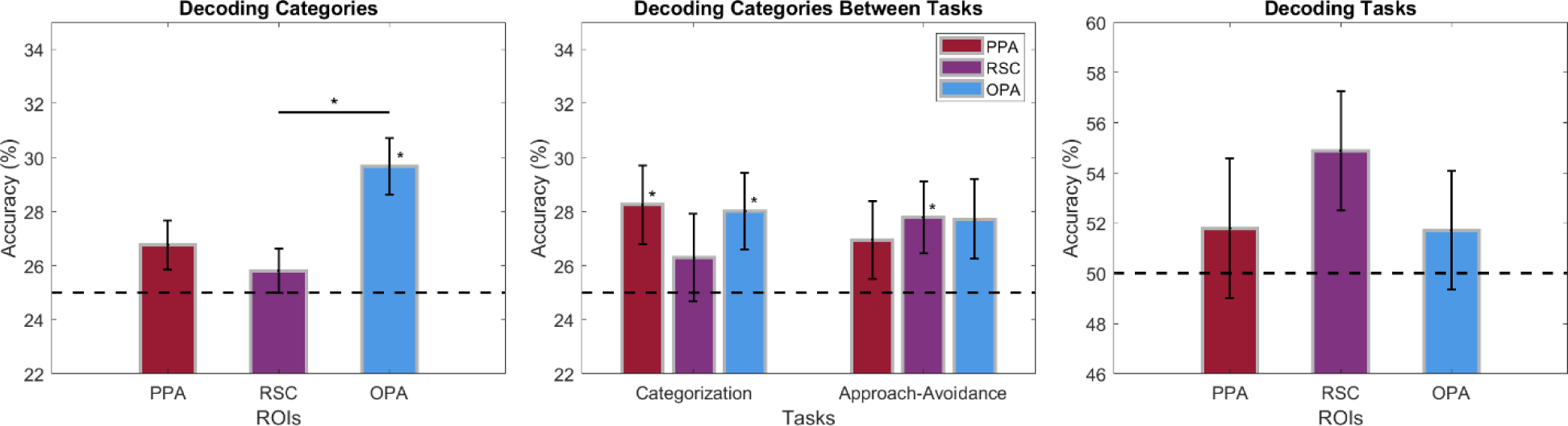
MVPA decoding results by ROI. Dashed lines indicate the chance levels. Significant decoding performances are indicated by asterisks right above the corresponding bar. Significant differences between ROIs are at p<0.05, Bonferroni corrected. Error bars represent ± 1 SEM.

Additionally, the one-sample t-tests applied to determine the significance of decoding performances compared to the chance level revealed several strong effects. The overall category decoding performance of the OPA was significantly above the chance level (p=0.0002). During the categorization task, category decoding performance was significantly above the chance level for the PPA (p=0.0371) and the OPA (p=0.0442). During the approach-avoidance task, only the RSC had a significant category decoding performance (p=0.0499). Finally, although none of the ROIs had significant task decoding performance, the RSC was close to the cut-off (p=0.0524).

### 3.3 RSA Results

We performed a model-based RSA defining responses to each stimulus under each task as separate conditions to examine the activity patterns in scene-selective regions compared to various candidate models. The resulting 64×64 neural RDMs and the MDS plots (metric stress minimized) can be seen in Fig. 6.

**Fig. 6.**
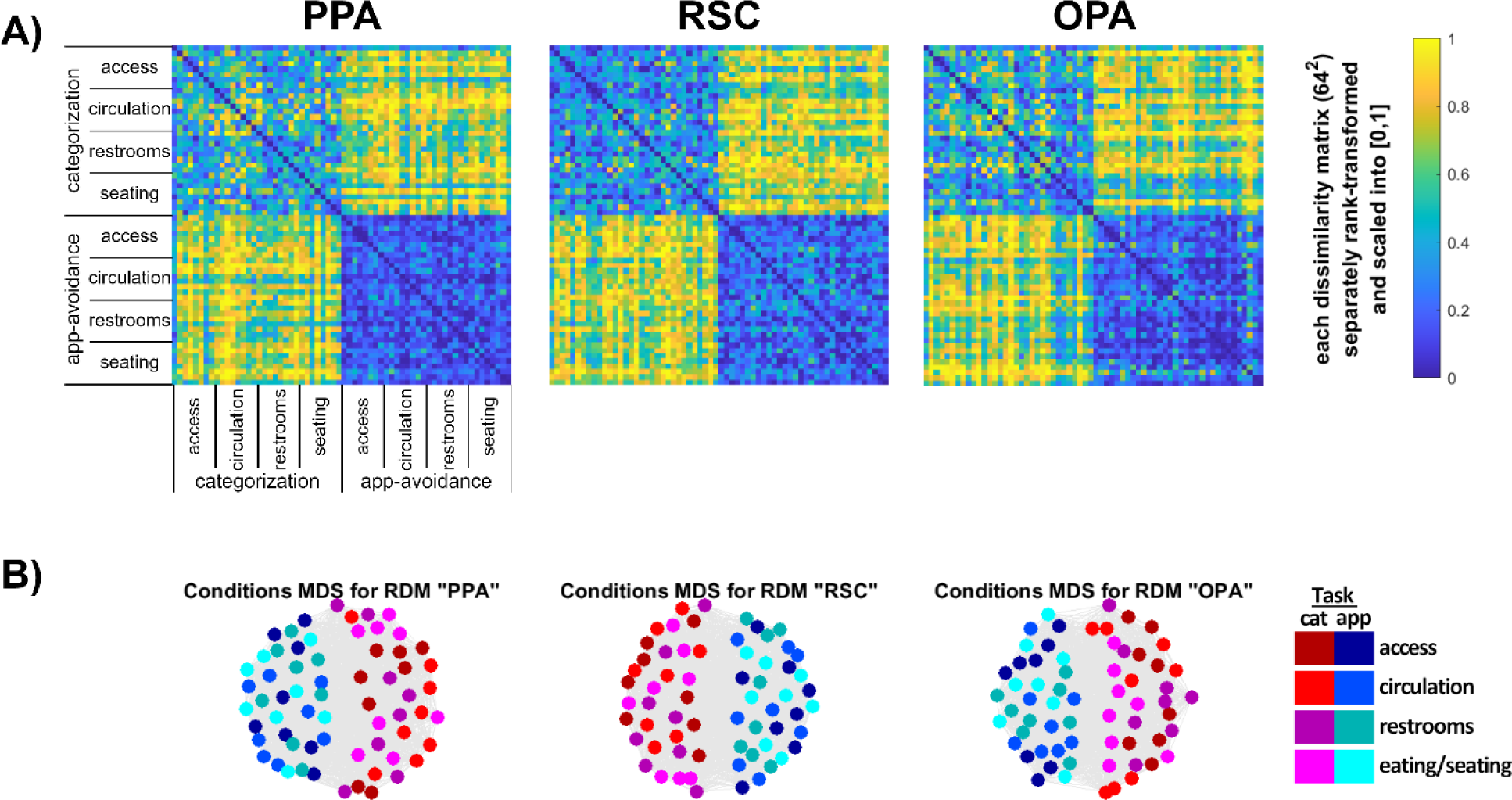
RSA results. (A) Neural RDMs for each ROI. Stimuli are represented by individual cells in the matrix, ordered by categories and tasks. (B) Conditions MDS plots for each ROI. The spatial distances between color-coded stimuli for each task are visualized.

These RDMs were then compared to the candidate models explained in section 2.7.4 by a signed-rank random-effects test corrected for multiple comparisons (FDR) at a threshold of p<0.01 via Kendall’s τ. As shown in Fig. 7A, the Task model was significantly similar to the neural RDMs for all ROIs. However, it did not reach the noise ceiling in any of these regions, indicating that it did not account for all the variance in the activation patterns. None of the other models were found to be similar to the neural RDMs.

**Fig. 7.**
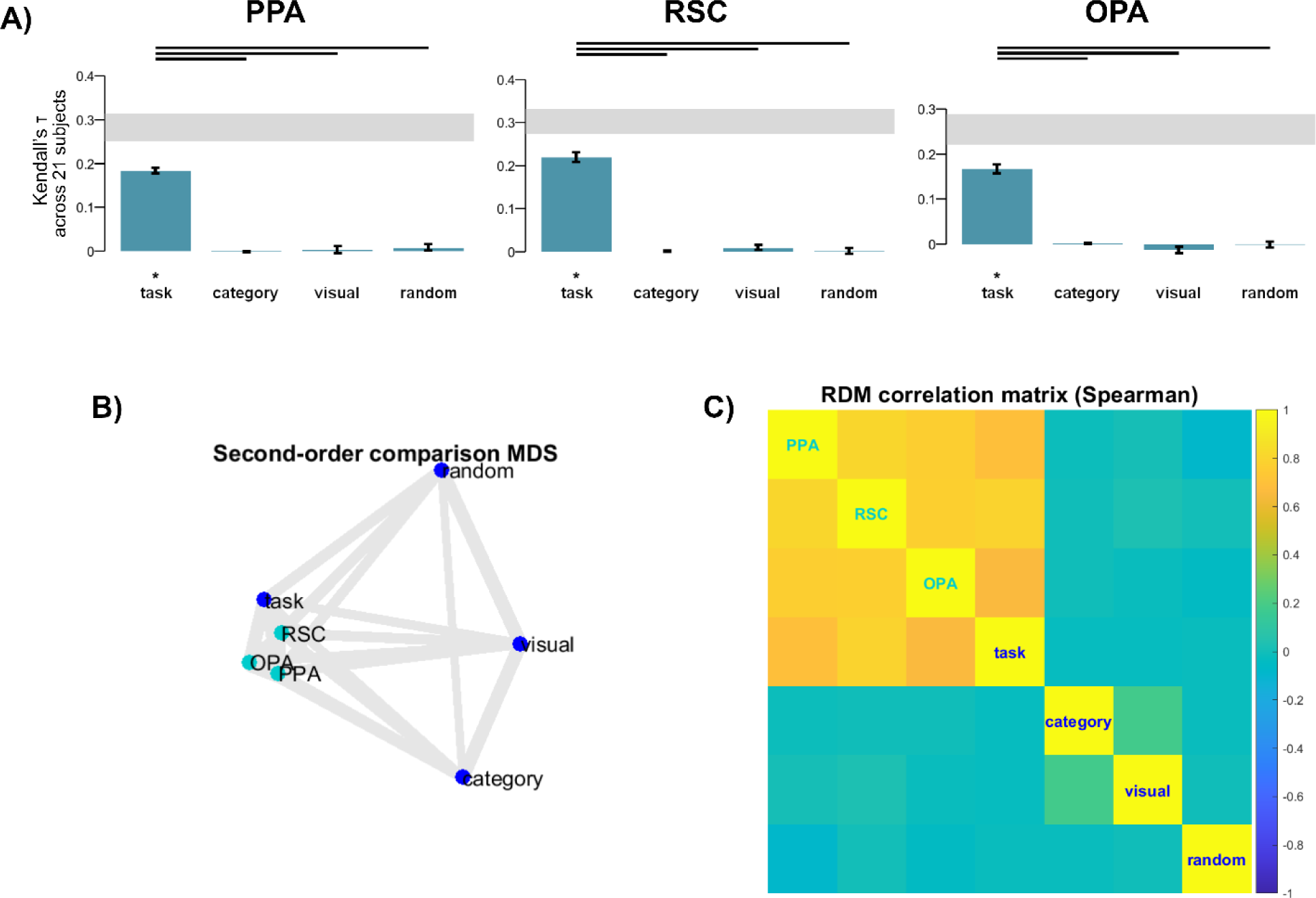
RDM-candidate model comparison results. (A) Kendall’s τ comparisons of ROIs to candidate models. The noise ceiling for a hypothetical true model is indicated with the gray bands over the graphs. The significant results are indicated with an asterisk at p<0.01. (B) Second-order comparison MDS, indicating the spatial distances between neural and candidate RDMS. (C) Spearman correlations of all the RDMS.

Precisely, Kendall’s τ values for the model comparisons were as follows. For PPA: task (τ=0.184, p=0.000), category (τ=-0.002, p=0.939), visual (τ=0.004, p=0.152), random (τ=0.008, p=0.144). The task model was significantly similar to the PPA but did not reach within the bounds of the noise ceiling (0.250, 0.314). The findings were similar for the RSC: task (τ=0.219, p=0.000), category (τ=-0.000, p=0.659), visual (τ=0.009, p=0.035), and random (τ=0.002, p=0.341). The task model was also significantly similar to the RSC but did not reach the noise ceiling (0.274,0.332). Finally, the same trend was observed for the OPA: task (τ=0.166, p=0.000), category (τ=0.001, p=0.293), visual (τ=-0.013, p=0.968), random (τ=-0.001, p=0.541). The task model was again significant, still not reaching the noise ceiling (0.221, 0.289).

In addition, second-order comparisons of all the neural and model RDMs revealed significant similarities between ROIs and the task model in general. At the same time, they all differed from other candidate models—the second-level comparison MDS can be seen in Fig. 7B and the Spearman correlation matrix of all the RDMs can be seen in Fig. 7C.

## 4. Discussion

This study used classification and RSA techniques to examine the effect of built environment categories and behavioral tasks on scene-selective regions in the human brain. The findings indicate that indoor categories were successfully decoded in the OPA in general. In contrast, the PPA and the OPA were significantly successful when we only looked at their performances during the categorization task, and the RSC was significantly successful only during the approach-avoidance task. The tasks were not successfully decoded in any of these regions. When we looked at the activation patterns to the categorical stimuli under each task and compared them to candidate task, category, visual, and random models, we observed that the task model was significantly similar to the actual neural activation patterns in all the ROIs. In contrast, other models were not at all significant. Despite being strongly correlated to the neural RDMs for all ROIs, the task model did not reach the noise ceiling, indicating that there are other factors a potential true model would incorporate that our models did not account for.

### 4.1 Built-environment categories are decoded best in the OPA, and the task affects decoding performance in all scene-selective regions

When we looked at the overall categorization performance of the ROIs, we see that although above chance in all the ROIs, it was only significantly high in the OPA. Also, the OPA was significantly more successful than RSC regarding category decoding performance. Although one would expect the PPA to have a significant category decoding performance based on the previous literature as it is pronounced in processing semantic information and identifying our surroundings (Dilks et al., 2022; Persichetti & Dilks, 2019), it was not observed in this analysis when we used data from both tasks in the classification process. This could result from a task effect, which will be discussed later. However, despite not being considered the most prominent in category discrimination, the overwhelming success of the OPA activity in decoding built environment categories can be explained by its proposed role in a particular type of navigation and how it may have been reflected in the criteria of our categorization approach, rather than it processing categorical information. Our categories of access points, circulation elements, restrooms, and eating/seating areas are essentially defined by their functions as part of a building, and they differ significantly by affordances. The access points include wide entrance areas, including the surroundings of buildings and doors one can walk through, and circulation elements consist of halls and stairs that primarily serve movement within a building. On the other hand, restrooms and eating/seating areas are less about movement and navigation, and these areas are less open and contain more borders, corners, and obstacles. The OPA may have been better at classifying these categories due to its involvement in *visually-guided navigation*, which concerns the immediate space visible to the subject, based on detecting paths and avoiding any obstacles by its sensitivity to borders, obstacles, and the openness of a scene (Dilks et al., 2022; Patai & Spiers, 2017; Persichetti & Dilks, 2018), and also its processing of navigational affordances even when they are not task-relevant (Bonner & Epstein, 2017, 2018).

Interestingly, when we analyzed the two tasks separately, the category decoding performances of these ROIs changed. Both the PPA and the OPA were significantly above the chance level during the categorization task when the participants actively considered the categorical properties of the stimuli. As explained before, the PPA is expected to have good decoding performance for categories; however, in this case, it was only apparent when the task was related, indicating an effect of behavioral goals. On the other hand, only the RSC was significantly above the chance level for category decoding during the approach-avoidance task. This could be related to the nature of the task, possibly engaging multiple processes such as considering oneself in relation to the scene (egocentric reference frames) and spatial memory, including retrieval of previously learned scenes from memory (Alexander et al., 2022; Miller et al., 2014; Mitchell et al., 2018; Vann et al., 2009). One explanation concerning spatial memory could be the repeated exposure of participants to the same stimuli during the experiment. Indeed, although participants were instructed to give approach-avoidance responses based on their initial opinions and not worry about being consistent throughout, some participants later indicated trying to remember their previous responses in subsequent runs. Another explanation may come from a possible role of the RSC in approach-avoidance processes, which, to the best of our knowledge, is unexplored in humans but is supported by animal studies on approach-avoidance learning (Todd et al., 2019). More research is necessary to clarify the RSC’s role in related processes.

Overall, the decoding results indicate that the category decoding performances of scene-selective regions are modulated by the behavioral goals of the observers.

### 4.2 Activation patterns in scene-selective regions are best explained by the task model

The model-based RSA results indicated the Task model best explained the activation patterns in scene-selective regions. Still, the similarity did not reach the noise ceiling, which a true model that explains the variation perfectly would. This means that, although the task model was significantly similar to neural activation patterns, it does not perfectly explain the data. We suggest that a combination of factors may better explain activation patterns in these regions than discrete models for task, category, and visual properties of the stimuli since brain regions do not necessarily process single, separate properties. Indeed, according to the literature, despite each being more pronounced in different functions, all three scene-selective regions are affected by various low- and high-level properties (Groen et al., 2017). Also, the functional differences among scene-selective regions that are demonstrated in the past literature, such as variations in category and visual property processing (Dilks et al., 2022; R. A. Epstein & Baker, 2019), were not observed with RSA, as all three regions resulted in highly similar RDMs. This may be due to the nature of the RSA and the way it probes the data as we observed these differences in the classification analyses, and it is natural for different types of analyses to uncover distinct aspects of the data. Therefore, comprehensive models, possibly separately for each ROI, are necessary to enlighten the differential nature of scene processing these regions.

Another possible reason we did not observe any ROI differences could be related to our method of ROI selection based on our localizer experiment. As explained earlier, we used group-level activation maps to define ROIs and kept ROI sizes equal while not omitting any participants. This caused us to be very liberal with the thresholds. As a result, we may have had ROIs that included surrounding regions in some participants with smaller ROIs and introduced variability into the activity patterns we examined with these analyses. There exist different approaches in the literature, some conservative, such as defining individual ROI masks and omitting participants who do not have clusters that reach a strict threshold. Both have advantages and disadvantages, possibly introducing different biases to the analyses, and there is no single correct way of defining ROIs using localizers (Friston et al., 2006; Saxe et al., 2006). Here, we chose not to be too strict by only looking at a few voxels with very high responses from just a few participants who have them. We tried to have a broader, generalizable approach without losing any valuable data.

### 4.3 Limitations

Although we aimed to address important gaps in the literature, using a systematic categorization method, realistic stimuli, and multiple tasks to study the perception of built environments, this study also has certain limitations.

Instead of professional, retouched photos or strictly controlled, unrealistic images, we aimed to create an ecologically valid stimulus set by choosing ordinary places from a database. While this is a plus from one point of view, it may have also posed certain limitations since we cannot control the low-level variables. It is known that scene-selective regions are affected by low-level features of the stimuli, which may have added variability to the neural activity patterns we were investigating. However, low-level features are relatively well-studied compared to higher-level processes, and it has been communicated in the field that it is time to view scene processing as it is: a combination of multiple processes using realistic stimuli and tasks (Malcolm et al., 2016). Also, we attempted to address this limitation by adding a visual similarity model to the RSA, which did not result in any significant similarity to the neural activation patterns.

Another limitation is related to the content of our stimulus set. Since we followed the categorization method we chose while deciding on the stimuli and were also limited by the content of the database, some of the images may have affected the participants’ behavioral responses and the resulting neural activation patterns. Various characteristics of the environment, such as geometry and aesthetics, can affect brain activity and approach-avoidance tendencies (Shemesh et al., 2021, 2022; Vartanian et al., 2013, 2015), and we indeed found that some stimuli elicited unbalanced responses (Supplementary Fig. 1). This can be alleviated by performing a norming study first, and deciding on neutral stimuli if a similar task is to be used and the topic of study is the valence-related judgments. In addition, approach-avoidance tasks often result in variable responses even when the stimuli are controlled for valence due to individual tendencies, and again, we observed considerable individual differences in the distribution of approach-avoidance responses (Supplementary Fig. 2). In this case, however, we were not interested in the valence of the stimuli and only aimed to measure the initial processing by employing an approach-avoidance task, responses to which are not entirely explained by beauty or pleasantness judgments.

### 4.4 Future directions

This study was a first step towards a more ecologically valid study of scene perception, and future studies can expand our understanding of this topic in several directions. This study only used one branch of one particular categorization approach, and future research can make use of other aspects of this model, use other existing models, or develop new strategies to include various types of scene categories based on different criteria. In this study, we used multiple tasks. However, this was only a first step, and it is necessary to develop and use various ecologically valid behavioral tasks that represent real-life scene perception and our actions within scenes engaging a wide range of cognitive processes. Such in-depth and high-level study of scene perception contributes to uncovering new information regarding the neural and behavioral mechanisms underlying scene perception. In the long run, such approaches can provide a scientific basis for fields such as environmental psychology and neuroarchitecture to aid environmental and architectural design processes, resulting in improved spaces that both serve human needs and promote cognitive functions.

## 5. Conclusion

This study examined the task influence on the neural activity patterns of scene-selective regions to the scenes from built environment categories. We utilized multivariate analysis techniques such as decoding and RSA. We found that the OPA had overall the highest decoding performance for built environment categories, and task-specific category decoding performances were different in all of these regions, with the PPA and the OPA being significantly successful during the categorization task, and the RSC being significantly successful during the approach-avoidance task. Moreover, with RSA, we found that behavioral goals significantly explain the changes in the neural activity patterns in scene-selective regions while not reaching a true model level. This study is a first step in examining the task influence on scene perception in depth, contributing to our understanding of high-level scene processing.

## Acknowledgments

The authors would like to thank Aslı Eroğlu and Ömer Rençbereli for their support in data collection.

## Data Availability

The data and code are available upon reasonable request from the first author.

